# Asymmetric secretion in budding yeast reinforces daughter cell identity

**DOI:** 10.1101/483388

**Authors:** Valerie N. Thomas, Eric L. Weiss, Jennifer L. Brace

## Abstract

Asymmetric segregation of cellular factors during cell division produces two cells with different identities. This asymmetry underlies cell fate decisions as well as the ability to self-renew. Asymmetric segregation of protein and RNA to the growing bud of *Saccharomyces cerevisiae* generates a daughter cell with features distinct from its mother. For example, asymmetric segregation of the transcription factor Ace2 to the newly formed daughter cell activates a gene expression program unique to daughters. Ace2 activates a cohort of genes, including degradative enzymes, which facilitate cell separation exclusively from the daughter. This asymmetric secretion leaves a characteristic ‘bud scar’ chitin ring on the mother. We sought to determine the sufficiency of Ace2 to define a daughter cell state by generating an *ACE2* allele which localizes to both mother and daughter nuclei. When Ace2 asymmetry is lost, Ace2 target gene transcription and translation occur in both mother and daughter cells. However, we find that mother cells retain bud scars and maintain asymmetric daughter-specific secretion of the wall degrading enzyme Cts1. These findings demonstrate that while mothers are competent to transcribe and translate Ace2 targets, additional intrinsic factors reinforce the daughter cell state.

**TOC Summary:** Asymmetric segregation of the transcription factor Ace2 drives daughter-specific cell separation after cytokinesis. Cells engineered to express Ace2 targets symmetrically produce the cell separation enzyme Cts1. However, secretion remains asymmetric suggesting other daughter-specific factors are required to reinforce the daughter cell identity.

## Introduction

Asymmetric cell division is a fundamentally important process for generating cellular diversity, resulting in two cells with different identities. Unequal inheritance of cellular components such as organelles, proteins, and RNA during cell division is the basis for asymmetric cell division. There are multiple well-understood mechanisms that describe how transcription, translation and localization are coordinated to ensure preferential segregation of cell fate determination factors (Knoblich, 2010; Li, 2013; Chen *et al.*, 2016). One mechanism to control cell fate is driven by asymmetric inheritance of regulators that alter gene expression. In *Drosophila melanogaster*, asymmetric localization of the transcription factor *prospero* to the basal cell cortex of the neuroblast, from where the ganglion mother cell (GMC) will emerge, ensures its preferential inheritance to the GMC. In the GMC, *prospero* enters the nucleus to repress neuroblast-specific genes and reinforces the GMC cell fate (Doe *et al.*, 1991; Knoblich *et al.*, 1995; Spana and Doe, 1995).

Asymmetric cell division has long been appreciated as a fundamental mechanism to generate the variety of cellular diversity required for the development of a multicellular organism from a single fertilized egg. In addition to being a key driver of cellular diversity during development, asymmetric cell division is crucial for tissue renewal and stem cell maintenance. Disruption of the balance between these two processes may lead to abnormal growth, over proliferation, tumorigenesis and cancer (Furthauer and Gonzalez-Gaitan, 2009; Gómez-López *et al.*, 2014). Therefore, understanding the mechanisms that generate and maintain asymmetry have important implications for developmental biology, stem cell biology as well as tumorigenesis.

This evolutionarily conserved method of division, while unequivocally essential for metazoan development, is also important in single-celled eukaryotes. In the budding yeast, *Saccharomyces cerevisiae*, the newly formed daughter cell establishes an environment distinct from its mother. Polarized trafficking of protein and RNA to the growing daughter cell prior to cell division allows daughters to create a unique state upon division (Amon, 1996; Bobola *et al.*, 1996; Sil and Herskowitz, 1996; Li, 2013; Yang *et al.*, 2015). The transcription factor Ace2 localizes specifically to the daughter nucleus and helps define the daughter state by activating a cohort of genes required for cell wall degradation following cytokinesis (Dohrmann *et al.*, 1992; O’Conallain *et al.*, 1999; Colman-Lerner *et al.*, 2001). One of Ace2’s targets, the chitinase Cts1, is secreted exclusively from the daughter side where it degrades chitin in the cell wall between mother and daughter (Kuranda and Robbins, 1991; Colman-Lerner *et al.*, 2001). The consequence of this asymmetry is the retention of a chitin ‘bud scar’, a defining feature of the mother cell (Barton, 1950; Bartholomew and Mittwer, 1953; Pringle, 1991).

Ace2’s nuclear localization is tightly regulated during the cell cycle to ensure Ace2 gene targets are activated only once in a cell’s lifetime (Weiss, 2012). Throughout most of the cell cycle, Ace2’s nuclear localization sequence (NLS) is phosphorylated by cyclin-dependent kinase (CDK), preventing Ace2 nuclear entry (O’Conallain *et al.*, 1999; Sbia *et al.*, 2008). Just prior to cell division, CDK activity drops and dephosphorylation of the NLS permits Ace2 nuclear entry (O’Conallain *et al.*, 1999; Mazanka and Weiss, 2010). However, due to Ace2’s NES, the protein is rapidly exported from the nucleus (Mazanka *et al.*, 2008). To generate asymmetry, the daughter-localized NDR/LATS kinase Cbk1 with its co-factor Mob2 phosphorylates and blocks Ace’s nuclear export sequence (NES) from interacting with export machinery (Weiss *et al.*, 2002; Jansen *et al.*, 2006; Mazanka *et al.*, 2008; Brace *et al.*, 2011). While mother-localized Ace2 can exit the nucleus, the daughter-localized pool cannot, generating a system to asymmetrically concentrate Ace2 in the daughter nucleus (Mazanka *et al.*, 2008).

We have previously shown that cell identity is important for Ace2’s nuclear localization. Ace2 does not accumulate in nuclei unless first entering the daughter cell (Mazanka *et al.*, 2008). However, whether Ace2 nuclear localization is sufficient to create a daughter-like environment in a mother cell has not been thoroughly examined. Therefore, we sought to better understand the functional significance of Ace2 mother-daughter asymmetry. We generated an allele of *ACE2* that no longer requires the asymmetrically localized kinase Cbk1 for nuclear localization. This allele disrupts the nuclear asymmetry of Ace2, resulting in localization to both mother and daughter nuclei. We find that although Ace2 targets are expressed symmetrically, cells maintain asymmetric secretion of Cts1. Our analysis elucidates that daughter-cell identity depends on more than just transcriptional driven processes controlled by Ace2. Instead, daughter cell identity requires additional intrinsic factors that reinforce the daughter cell state.

## Results

### Mimicking constitutive Cbk1 phosphorylation breaks Ace2 asymmetry

Asymmetric Ace2 nuclear localization contributes to daughter cell identity in budding yeast. However, the sufficiency of Ace2 to establish this daughter state is currently unknown. We therefore sought to determine if mother cells would acquire a ‘daughter-like’ state if Ace2 were competent to accumulate in the mother nucleus. The kinase, Cbk1, directly controls the asymmetric nuclear accumulation of Ace2 via phosphorylation of two sites in the Ace2 nuclear export sequence (NES), preventing nuclear exit (Mazanka *et al.*, 2008). A third site in the C-terminus of Ace2 increases the expression of Ace2 target genes (McBride *et al.*, 1999; Mazanka *et al.*, 2008). We reasoned that constitutive phosphorylation of these sites would be sufficient to break Ace2’s asymmetry, expecting nuclear accumulation of Ace2 and activation of Ace2 target genes in both mother and daughter cell.

To this end, we mimicked constitutive Cbk1 phosphorylation by mutating Cbk1 phosphorylation sites to aspartic acid (S122D, S137D, and S436D, referred to as *ACE2^3D^*, Figure 1A). Expression of Ace2^3D^ was similar to wild type (Figure 1B) and cells exhibited a wild type cell separation phenotype (Figure 1C). To examine Ace2’s nuclear localization, we first distinguished mothers from daughters by briefly incubating cells with the cell wall-binding lectin concanavalin A (ConA) conjugated to rhodamine. Stained cells were then chased in label-free media for 3 hours to generate a population of stained mothers and unstained daughters (Mazanka *et al.*, 2008). We found GFP-tagged *Ace2^3D^* was nuclear enriched and accumulated in both mother and daughter nuclei (Figure 1, D-G). In 91% of large-budded wild type cells, Ace2 was found exclusively in the daughter nucleus with a small percent also exhibiting weak localization to mother nuclei (Figure 1E). Comparatively, in 90.3% of mother-daughter pairs, Ace2^3D^ localized to both the mother and daughter nuclei with only a few cells exhibiting asymmetry (Figure 1E).

**Figure 1.**
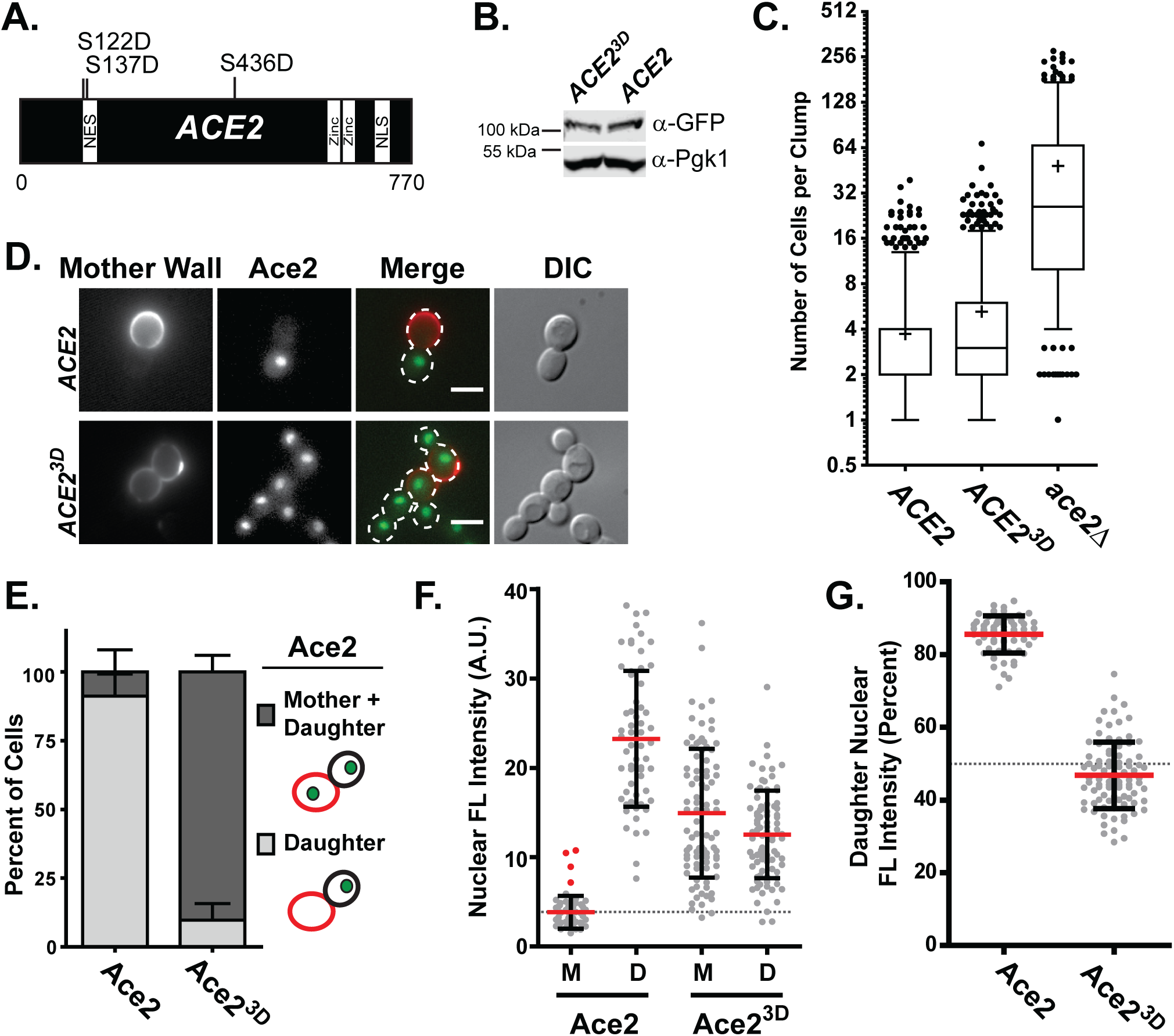
Mimicking Cbk1 phosphorylation in Ace2’s NES disrupts its asymmetric localization. (A) Schematic of Ace2 illustrating sites phosphorylated by Cbk1 and mutated to generate the ACE2^3D^ allele. NES = nuclear export sequence, Zinc = zinc-finger, NLS = nuclear localization sequence. (B) Ace2 protein levels are similar in ACE2 and ACE2^3D^ cells. Lysate from asynchronous cells expressing Ace2-GFP or Ace2^3D^-GFP were probed using a GFP antibody. A representative Western blot against GFP and Pgk1 (as loading control) is shown. (C) Ace2^3D^ is functional. Cell separation was quantified by counting the number of connected cells in a clump. A box and whisker (5 to 95 percentile) graph is shown. ‘+’ indicates the mean clump size. (D) Upon mutation of Cbk1 phosphorylation sites, Ace2 symmetrically localizes to mother and daughter nuclei. Representative max projection images of asynchronous cells expressing Ace2-GFP or Ace2^3D^-GFP. Mother cells are marked with rhodamine-concanavalin A (see methods). Scale bar is 5 microns. (E) Quantification of symmetric nuclear localization. The percent of large-budded cells exhibiting GFP localization to the ‘daughter’ or ‘mother and daughter’ nuclei is shown. No cell exhibited GFP signal in a mother nucleus only. The mean percentage and standard deviation of 3 independent experiments are shown (ACE2 n=38, 13, 19 cells; ACE2^3D^ n=35, 25, 42 cells). (F) Quantification of nuclear intensity. GFP intensity from mother and daughter nuclei was measured. In wild type mother cells with no nuclear enrichment, a region in the cytoplasm was measured (average, dashed grey line). Fluorescence (FL) intensity of each nuclei is plotted with standard deviation (mean, red line) from ACE2 n = 61 cells and ACE2^3D^ n = 89 cells (pooled from 3 independent experiments). (G) Total nuclear signal from mother-daughter pairs was calculated from nuclei in (F). The percent of the signal from the daughter nuclei is plotted with standard deviation (mean, red line). The grey line indicates 50% (equal mother-daughter signal).

To quantify the extent of Ace2 nuclear accumulation, we measured mother and daughter nuclear fluorescence intensity of GFP-tagged wild type Ace2 or *Ace2^3D^*. As demonstrated previously, wild type Ace2 demonstrated significant enrichment in the daughter nucleus with a few cells exhibiting weak nuclear signal in the mother (red dots) over the cytoplasmic background (grey line, Figure 1F; Colman-Lerner et al., 2001; Weiss et al., 2002). In contrast, the mean Ace2^3D^ nuclear intensity was elevated in mother nuclei relative to the weak signal in wild type mother nuclei and above the cytoplasmic background (Figure 1F). Consistent with the division of total Ace2 into two nuclei (Mazanka and Weiss, 2010), we found the mean fluorescence intensity of nuclear Ace2^3D^ in mother or daughter nuclei was lower than that of wild type daughter cells (Figure 1F). To confirm this, we calculated the percent of the total signal coming from the daughter. We found Ace2^3D^ nuclear accumulation to be approximately equal in the two nuclei (Figure 1G). Taken together, mimicking constitutive Cbk1 phosphorylation on Ace2 was sufficient to disrupt asymmetric Ace2 nuclear localization.

### Symmetric Ace2 localization is not sufficient to disrupt bud scar formation

Previous work demonstrated that disruption of Ace2’s NES is sufficient to drive symmetric localization of Ace2 (Colman-Lerner *et al.*, 2001; Sbia *et al.*, 2008). However, the function of Ace2 in the mother has not been fully characterized in these cells. In daughter cells, Ace2 activates a cohort of daughter-specific genes, the products of which are secreted to degrade chitin in the cell wall between mother and daughter cell at the end of cell division (Kuranda and Robbins, 1991; Colman-Lerner *et al.*, 2001; Doolin *et al.*, 2001; Baladrón *et al.*, 2002; Weiss *et al.*, 2002). Residual chitin on the mother cell, which is not degraded due to limited diffusion of chitinase, results in the formation of a bud scar, which is commonly used to identify mother cells (Barton, 1950; Pringle, 1991). We reasoned if Ace2 activation occurs in both mother and daughter cell, symmetric degradation of chitin from both sides of the cell would eliminate the bud scar. To test whether *ACE2^3D^* mother cells retain bud scars, we labeled mother cells with rhodamine-ConA and chased in label-free media for 2 hours. Then residual chitin in the bud scar was stained with the chitin-binding dye calcofluor white. As expected, most wild type mother cells had between 1 and 3 bud scars, with a smaller percentage having 4 or more (Figure 2, A and B). Additionally, a small number of unstained daughter cells exhibited the presence of a bud scar. This likely represents the chitin ring that is synthesized in the daughter cell just prior to bud emergence, which is indistinguishable from a bud scar in this assay (Figure 2A asterisk, (Shaw *et al.*, 1991)).

**Figure 2.**
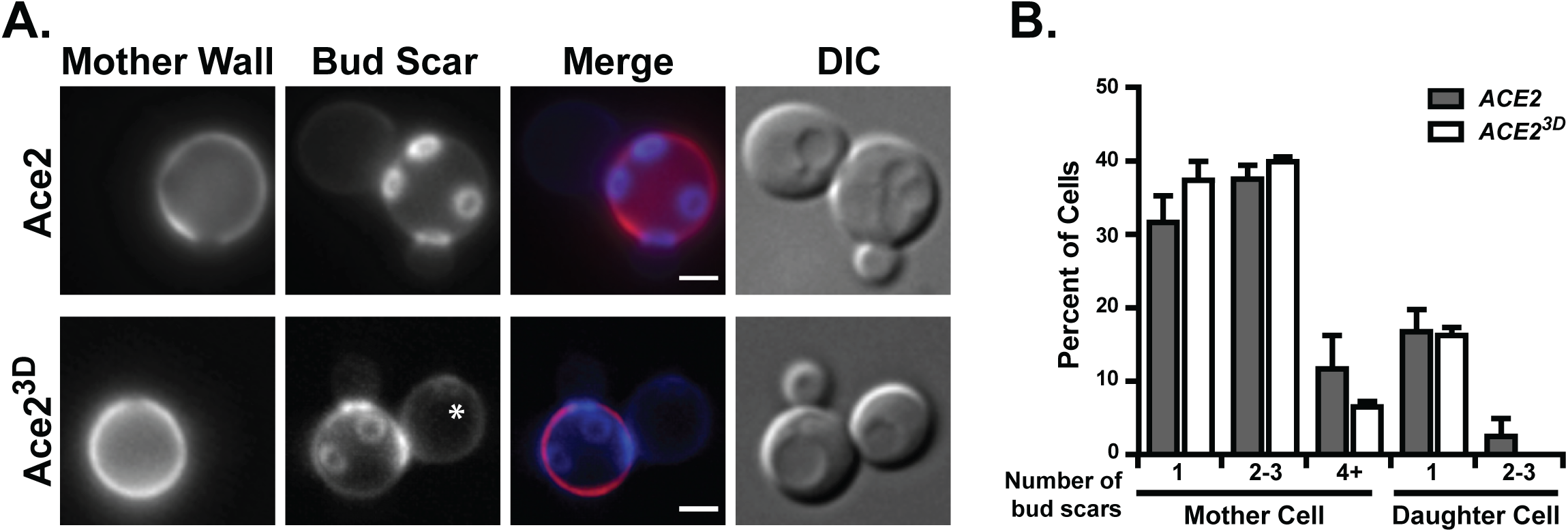
Bud scar asymmetry is maintained in cells with symmetric nuclear Ace2. (A) Ace2^3D^ expressing cells maintain mother-specific bud scars. Shown are representative max projection images of asynchronous cells of *ACE2* or *ACE2^3D^* genotype. Mother cells are labeled with concanavalin A (red) and bud scars are visualized using calcofluor white (blue). An example of a daughter chitin ring forming at bud emergence is marked with an asterisk (*). DIC (differential interference contrast) shows cell outline. Scale bar is 2 microns. (B) No difference in the presence or quantity of bud scars between cells expressing *ACE2* or *ACE2^3D^*. Plotted is the mean percent of cells with the indicated number of bud scars present on the mother or daughter cell. The standard deviation of 3 independent experiments (*ACE2* n=46, 28, 45 and *ACE2^3D^* n=57, 30, 39) is shown. A two-tailed equal variance student’s t-test was used to determine that there was no significant difference between *ACE2* and *ACE2^3D^* (p > 0.05) for each bud scar category.

More importantly, we found Ace2^3D^ mother cells retained bud scars (Figure 2, A and B). Similar to wild type, *ACE2^3D^* mother cells had between 1-3 bud scars with a small percentage having 4 or more (Figure 2B). These data were surprising and suggested disrupting Ace2’s nuclear asymmetry was not sufficient to disrupt asymmetry in processes downstream from Ace2.

### Ace2^3D^ exhibits transcriptional activity in mother cells

Retention of bud scar asymmetry, despite Ace2 nuclear symmetry, could be explained by a lack of Ace2-target gene expression in mother cells. To determine if the *ACE2^3D^* allele can activate transcription, we first measured the transcript levels of a variety of Ace2 targets (*CTS1*, *SCW11*, *DSE1*, *DSE2*, *DSE3* and *DSE4*) by quantitative PCR (qPCR) in asynchronous cultures (Figure 3A). Relative to the wild type strain, we found Ace2 target transcripts were either produced at levels similar to wild type (*CTS1*, *SCW11*, *DSE4*) or were slightly elevated compared to wild type (*DSE1*, *DSE2*, and *DSE3*). While these data demonstrate the ability of Ace2^3D^ to activate transcription, it does not allow us to discriminate activity in mother versus daughter cells.

**Figure 3.**
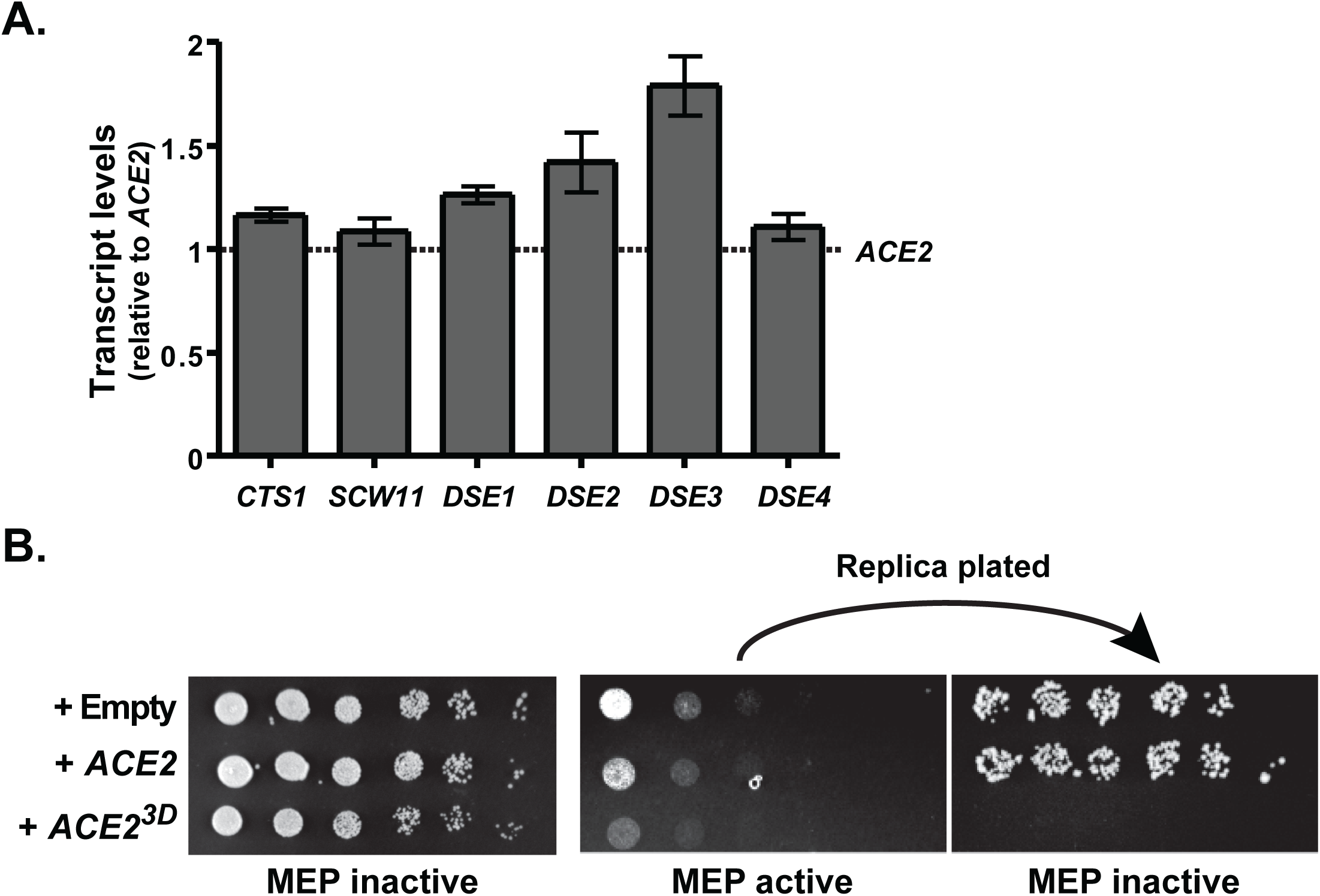
Ace2 target promoters are activated in mother cells expressing *ACE2^3D^*. (A) Ace2 target transcripts are comparable between *ACE2* and *ACE2^3D^* cells. Transcripts were measured by real-time quantitative PCR and normalized to actin (*ACT1*) levels. The average fold change in expression in an asynchronous culture relative to the *ACE2* control strain is plotted. Error bars represent the standard deviation of 3 independent experiments. (B) Mother Enrichment Program (MEP) cells expressing *ACE2^3D^* do not recover from estradiol-induced cell death. MEP cells transformed with a plasmid containing an empty vector, *ACE2* or *ACE2^3D^* were plated to plates containing estradiol (MEP active) or vehicle control (MEP inactive). After 3 days of growth, cells were replica plated from MEP active plates to plates without estradiol (MEP inactive) to allow mother cells that survived to propagate daughters.

To more directly demonstrate Ace2 transcriptional activity in mother cells, we exploited a genetically engineered strain termed the mother enrichment program (MEP). The MEP strain utilizes the daughter-specific activity of Ace2 to enrich for mother cells through an Ace2-target-mediated cell death (Lindstrom and Gottschling, 2009). In this system, an Ace2 regulated promoter drives expression of Cre recombinase in a strain with two essential genes flanked by LoxP sites. In wild type cells, Ace2-driven Cre kills daughter cells, leaving viable mother cells. To permit strain growth, Cre is post-transcriptionally regulated by fusion to an estradiol-binding domain allowing nuclear localization of Cre only in the presence of estradiol.

We reasoned that if Ace2^3D^ activates target genes in mother and daughter cells, then both mother and daughter would be killed in the MEP strain. To test this, we transformed the MEP strain with a plasmid expressing *ACE2* or *ACE2^3D^* from its endogenous promoter. Cells were propagated in the absence of estradiol and spotted to media containing estradiol to activate the MEP (MEP active) or an ethanol control plate (MEP inactive). MEP cells containing the empty vector control or expressing *ACE2* exhibited little growth on plates containing estradiol. Mother cells increase in size but daughter cells do not propagate (Figure 3B). Interestingly, cells expressing the *ACE2^3D^* allele exhibited no background growth, suggesting mothers were also killed. To confirm this lethality, cells exposed to estradiol were replica plated to vehicle control plates to allow outgrowth of surviving cells. MEP cells with empty vector or expressing *ACE2* recovered, but cells expressing *ACE2^3D^* did not (Figure 3B, replica plated). This result suggests that Ace2^3D^ can drive expression of its transcriptional targets in both mother and daughter cells.

### Mothers expressing Ace2^3D^ translate Ace2 targets

Given Ace2 target transcription occurs in mother cells, the retention of bud scars on mothers could be explained by an inability of mothers to translate messages produced by Ace2. Therefore, to unambiguously distinguish expression in mothers from daughters, we used live-cell imaging to examine the localization of Ace2 targets. In wild type cells, Ace2-driven *CTS1* transcripts are translated into the lumen of the endoplasmic reticulum (ER), trafficked through the endomembrane system to the daughter bud neck, and then secreted outside the cell. Cts1 has an ER-targeting signal sequence (SS) at its N-terminus, followed by a catalytic domain, a serine/threonine rich (S/T rich) glycosylation region and a chitin binding domain (CBD) (Figure 4A, (Hurtado-Guerrero and van Aalten, 2007)). To minimize disruption to endogenous Cts1 expression, we tagged Cts1 internal to the coding sequence with a monomeric superfolding GFP (msGFP), which folds properly in the endomembrane system (Fitzgerald and Glick, 2014; Day *et al.*, 2018). This methodology leaves the 5’ and 3’ regulatory elements intact (Dohrmann *et al.*, 1996; Aulds *et al.*, 2012; Wanless *et al.*, 2014). To avoid aberrant vacuole targeting of the msGFP tag, we also introduced the *vps10-104* allele (Fitzgerald and Glick, 2014). Additionally, we found that media pH altered the population of Cts1 we could visualize. Under acidic conditions (minimal media), Cts1 was visible in cellular internal structures like the ER and cytoplasmic puncta, while in neutral media (rich media or buffered minimal media) we observed bud neck localization likely representing the secreted or extracellular Cts1 (Figure S1). Since the signal intensity of the secreted material masked the weaker internal signal, most examination of Cts1 localization was performed in minimal acidic media.

**Figure 4.**
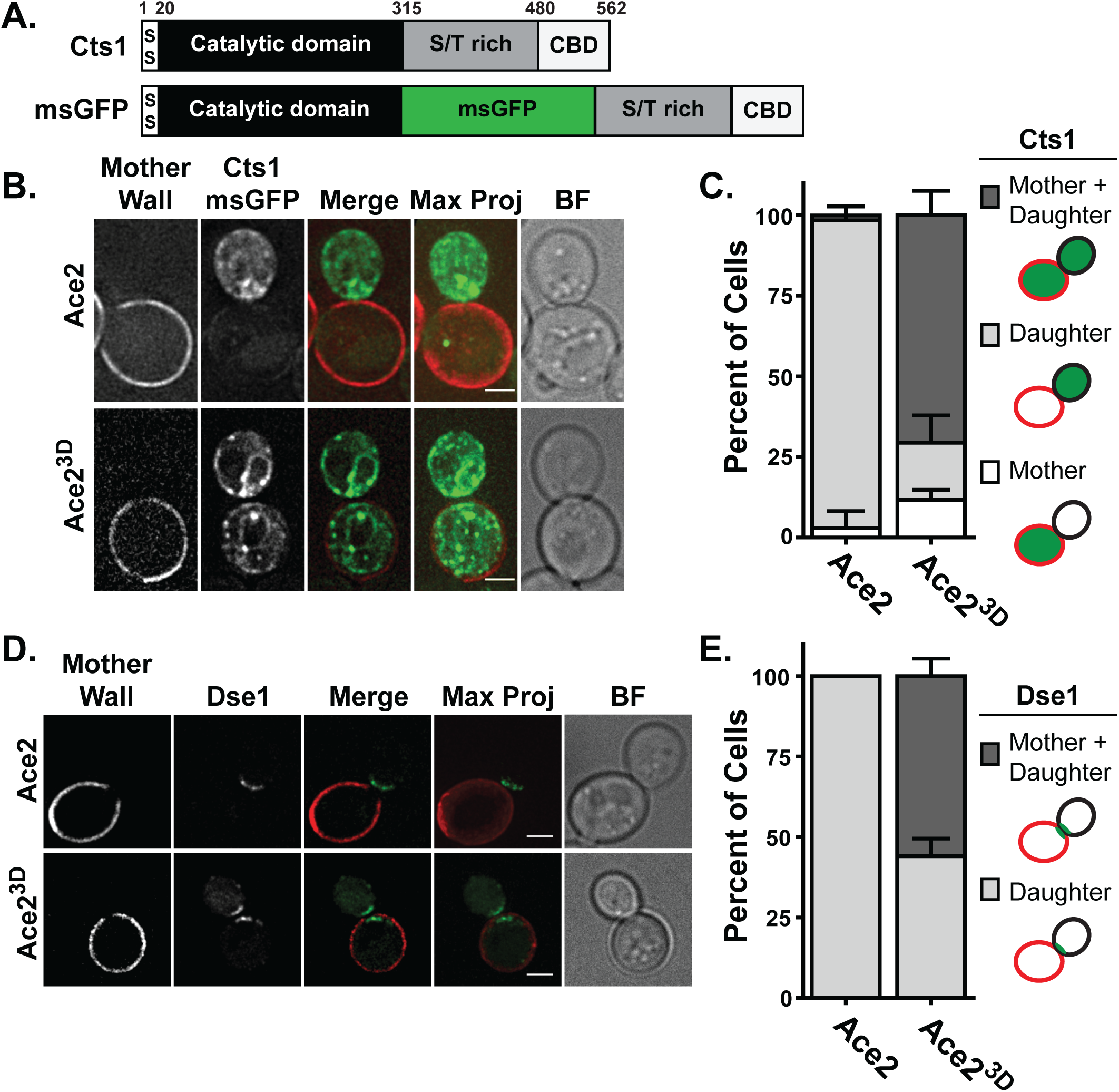
Mother cells expressing *ACE2^3D^* translate Ace2 targets. (A) Schematic of Cts1 and msGFP-tagged Cts1. SS, signal sequence; S/T rich, serine/threonine rich glycosylation region; CBD, chitin binding domain; msGFP, monomeric superfolder GFP. (B) ACE2^3D^ disrupts daughter-specific Cts1 translation. Shown are representative images of *ACE2 vps10-104* or *ACE2^3D^ vps10-104* cells expressing the Ace2 target Cts1 tagged with msGFP (green). Mother cells are marked with concanavalin A (red). BF (brightfield) allows visualization of the cell outline. Scale bar is 2 microns. See also Figure S2. (C) *ACE2^3D^* cells quantitatively exhibit Cts1-msGFP expression in both the mother and daughter cell. In large-budded, mother-daughter pairs from (A), the mean percent of cells with Cts1-msGFP signal in both mother-daughter (dark grey bar), daughter only (light grey bar) or mother only (white bar) are shown. Error bars represent standard deviation of 3 independent experiments (*ACE2* n=22, 11, 21; *ACE2^3D^* n=17, 33, 28). (D) The Ace2 target Dse1 expression in mother cells with the *ACE2^3D^* allele. Representative images of asynchronous *ACE2* or *ACE2^3D^* cells expressing Dse1-GFP (green). Mother cells are marked with concanavalin A (red), and BF (brightfield) shows cell outline. Scale bar is 2 microns. (E) Quantification of ACE2^3D^ cells exhibiting Dse1-GFP expression in both the mother and daughter cell. The percent of mother-daughter pairs expressing Dse1 in both mother and daughter (dark grey bar) or daughters only (light grey bar) is shown. No cases of mother only expression were observed. Error bars represent standard deviation of 3 independent experiments (*ACE2* n=49, 68, 48 cells; *ACE2^3D^* n=40, 34, 55 cells).

In wild type cells, Cts1 protein was observed almost exclusively in the daughter cell and could be seen in the ER and in cytoplasmic puncta (Figure 4, B and C). In contrast, both mother and daughter cells exhibited Cts1 localization to the ER and cytosolic puncta in the *ACE2^3D^* strain. To quantify this, we analyzed large-budded, mother-daughter pairs and found a significant increase in the percent of pairs in which Cts1 was localized to both mother and daughter (70.6% *ACE2^3D^* vs 1.6% *ACE2*, *p*=0.0001 two-tailed, equal variance student’s t-test, Figure 4C).

As an additional indication that mother cells can translate Ace2 targets, we isolated a serendipitous mutant while tagging the endogenous locus with msGFP. A spontaneous nucleotide deletion generated a truncation allele of Cts1 (Cts1^trunc^) lacking the majority of its C-terminus including the serine-threonine glycosylation region (Figure S2A). This mutant fails to exit the ER and is clearly not expressed in, or excluded from, the mother in wild type cells (Figure S2B). In contrast, *ACE2^3D^* cells exhibit ER localization of Cts1^trunc^ to both mother and daughter, further supporting our observations that mothers express Cts1.

To further analyze expression of Ace2 targets in mother cells, we examined another Ace2 target, one that does not transit the endomembrane system. The Ace2 target Dse1 localizes to the daughter bud neck in large budded cells (Frýdlová *et al.*, 2009). We C-terminally tagged the endogenous Dse1 locus with GFP and examined its localization in mother-daughter stained cells. Similar to Cts1, Dse1 was found exclusively in the daughter cell in wild type cells (Figure 4, D and E). Conversely, Dse1 was clearly localized to the mother cell in *ACE2^3D^* cells. On average, 56% of mother-daughter pairs exhibited localization to both mother and daughter while no wild type cells exhibited this localization pattern (p < 0.0001, two-tailed, equal variance student’s t-test). Taken together, these data suggest that transcripts produced via Ace2^3D^ in mother cells are likely translated in mother cells.

### *ACE2^3D^* expressing cells maintain asymmetric secretion of chitinase

Since transcription, translation and localization of Ace2 targets occurs in *ACE2^3D^* mother cells, we next investigated the possibility that the presence of bud scars in these cells could be explained by the inability of mothers to secrete Cts1. To directly visualize Cts1 secretion, we examined the external population of Cts1 by imaging in buffered minimal media. The C-terminus of Cts1 contains a chitin binding domain (CBD), which is thought to retain chitinase in the bud neck to promote efficient cell separation (Hurtado-Guerrero and van Aalten, 2007). Unfortunately, this caused preferential retention of Cts1 to the mother side chitin, masking our ability to assess the origin of secretion. Therefore, we expressed an allele of Cts1 lacking the CBD (Cts1 ΔCBD, Figure 5A). We found Cts1 ΔCBD was no longer retained in bud scars and only exhibited transient signal at the bud neck, allowing us to image the recently secreted Cts1. To mark the plasma membrane (PM), we used a mCherry tagged plextrin-homology domain and mothers were identified by bud scar staining or size (see methods). Line scan analysis across the bud neck from mother to daughter in wild type cells illustrated that peak Cts1 signal occurred more proximal to the daughter plasma membrane (Figure 5, A and B). We observed the same proximity of Cts1 signal to the daughter PM in cells expressing *ACE2^3D^*. Quantification of the distance between peak Cts1 signal to peak mother or daughter PM demonstrates, on average, a shorter distance to the daughter PM than the mother PM in both genotypes

**Figure 5.**
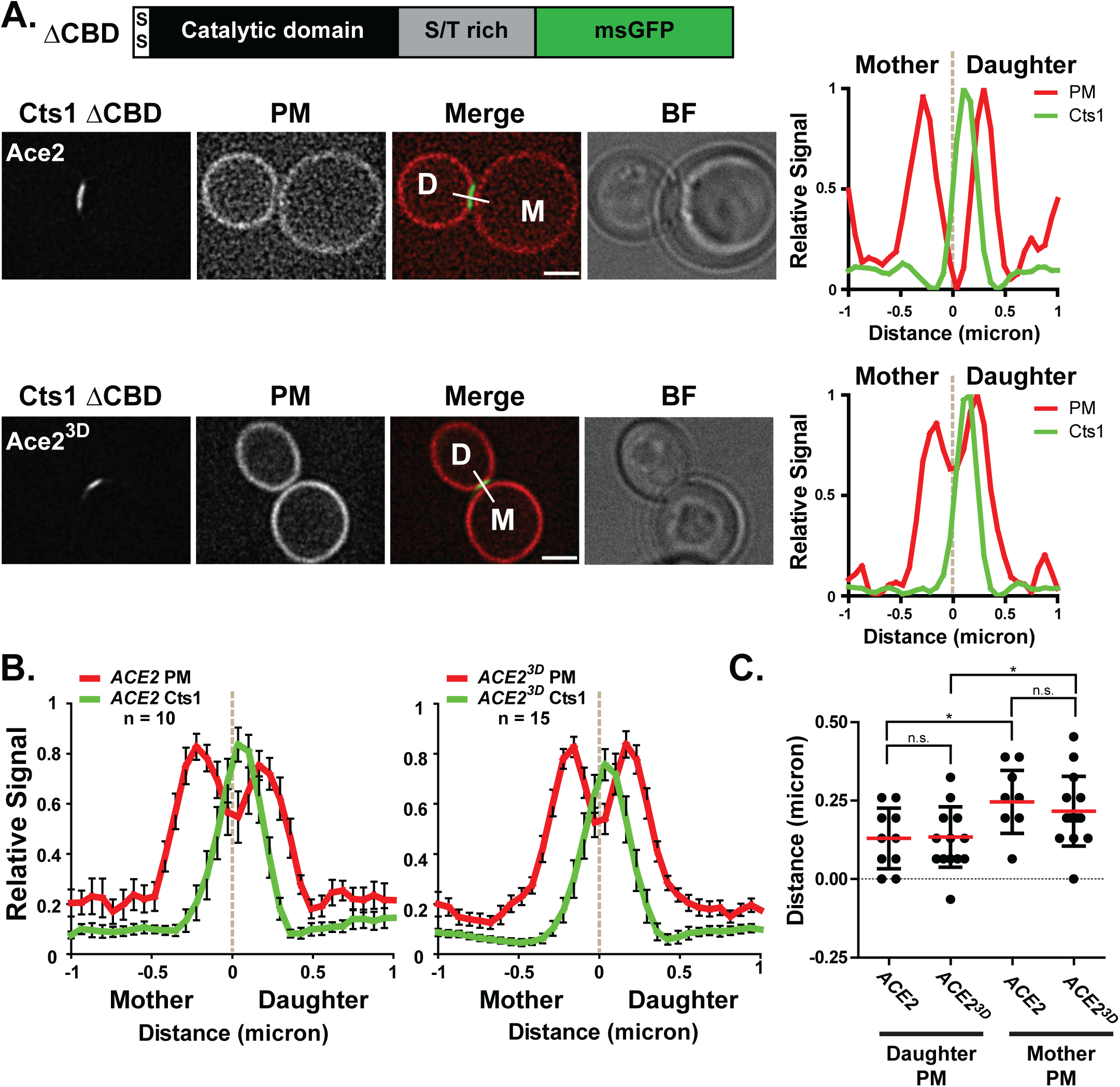
Secretion of the Ace2 target, Cts1, occurs exclusively from the daughter cell. (A) Cts1 is secreted preferentially from the daughter side, despite expression in both mothers and daughters in *ACE2^3D^* cells. A schematic of the Cts1 ΔCBD-msGFP is shown above a representative single-slice image of asynchronous *vps10-104 ACE2* or *ACE2^3D^*cells expressing Cts1 ΔCBD-msGFP (green). The plasma membrane (PM) is marked with PLC-delta PH domain fused to mCherry (red). BF (brightfield) microscopy shows cell outline. Scale bar is 2 microns. To the right of the image, a plot of the signal intensity for msGFP and mCherry across the line drawn at the site of secretion from mother to daughter is shown. Peak intensity for each channel was scaled between 0 and 1 and the line’s center, between the PM peaks, was set to 0 and is indicated by the vertical grey dashed line. (B) Secreted Cts1 is skewed to the daughter side regardless of genotype. A plot of the average signal intensity of GFP and mCherry across a line drawn at the bud neck as in (A) is shown. Error bars represent the standard error of the mean (*ACE2* n=10; *ACE2^3D^*n=15). (C) The peak of secreted Cts1 occurs closest to the daughter cell plasma membrane. For each cell analyzed in (B), the distance of the Cts1 peak to the daughter PM and the mother PM was determined and plotted. The average distance is shown in red with standard deviation. p-values are as follows: * = 0.01 to 0.05, n.s. > 0.05 (two-tailed, unpaired student’s t-test).

(Figure 5C). Thus, we find that although mother cells are capable of expressing Ace2 targets, additional mechanisms ensure mothers do not secrete Cts1.

## Discussion

In this study, we sought to understand the functional significance of asymmetric cell division in *S. cerevisiae* by disrupting the asymmetry of a key daughter-specific factor. We found that simply disrupting the asymmetric localization of the daughter-specific transcription factor Ace2 was not sufficient to induce a daughter state in the mother cell. Despite symmetric expression of the cell wall degrading enzyme Cts1, cells maintained asymmetric secretion of Cts1 and degradation of the cell wall from the daughter cell. These results suggest that more complex, daughter-specific mechanisms likely work together to ensure asymmetric cell division in *S. cerevisiae*.

### Ace2 transcriptional asymmetry is not sufficient to control cell fate

The transcription factor Ace2 is one of several asymmetries in budding yeast (Amon, 1996; Li, 2013; Yang *et al.*, 2015). Exquisite control of Ace2 during the cell cycle ensures that it is activated only once in a cell’s lifetime (Weiss, 2012), suggesting it is a major player in controlling cell identity. Similar to an established mutation in Ace2’s NES (G128E) (Sbia *et al.*, 2008), we demonstrate here that mutations mimicking constitutive Cbk1 phosphorylation (*ACE2^3D^*) are sufficient to disrupt Ace2’s asymmetric nuclear localization. We find, taking advantage of a previously established strain which kills cells that activate Ace2 transcription, that the *ACE2^3D^* allele kills both mother and daughter cells, providing strong evidence that this allele activates target gene transcription from both the mother and daughter cell.

Nevertheless, our data suggests that Ace2 alone is not sufficient to induce a daughter cell state, as mother cells retain bud scars. To rule out the possibility that Ace2 target transcripts are not translated in mother cells, we examined the localization of an Ace2 target, Cts1, in live cells. Previous work to examine Cts1 localization used C-terminal tagging of Cts1 with fluorescent proteins (Colman-Lerner *et al.*, 2001; Ríos Muñoz *et al.*, 2003) or the Cts1 promoter was used to drive expression of fluorescent proteins (Colman-Lerner *et al.*, 2001). However, in the oxidative environment of the ER, GFP folding is compromised leading to aberrant oligomers and reduced fluorescence (Aronson *et al.*, 2011; Costantini and Snapp, 2013). To circumvent these issues, we tagged Cts1 with a monomeric superfolder GFP (msGFP) variant that has been used to successfully image the lumen of the ER in yeast (Fitzgerald and Glick, 2014). When Ace2 is symmetrically localized, we clearly observe Cts1-msGFP signal in the ER and cytoplasmic puncta in both mother and daughter. This is probably not due to diffusion from daughter into the mother cell, as wild type cells demonstrate distinct expression in daughters only. Interestingly, even a Cts1 truncation mutant that fails to exit the ER exhibits exquisite daughter-specific localization, which is disrupted in *ACE2^3D^* expressing cells. Given that ER luminal, but not membrane-bound, proteins are able to freely diffuse between mother and daughter (Luedeke *et al.*, 2005), it is possible that Cts1 may be tethered in the daughter cell by a membrane-spanning protein that restricts its diffusion into the mother cell. Alternatively, Cts1 translation may occur after cytokinesis, once the ER lumen is physically separated. In either case, we find Cts1 transcripts produced in the mother are likely also translated in the mother.

### Additional daughter-specific mechanisms ensure daughter cell fate

Despite expression of Ace2 targets in mother cells, *ACE2^3D^* mother cells do not secrete Cts1, suggesting Cts1 secretion is dependent on the daughter cell state. How does Cts1 secretion remain asymmetric upon loss of its asymmetric translation? One possibility is that secreted cargos are trafficked differently depending on where translation occurred. In higher eukaryotes, evidence is building that asymmetry in the endomembrane system plays an important role in cell fate and polarity. For example, during neuron morphogenesis, spatial organization and polarization of Golgi outposts promote asymmetric dendrite growth through the polarized delivery of cargos (Horton *et al.*, 2005). Additionally, in fly sensory organ precursor cells, asymmetric inheritance of endosomes contributes to cell fate (Coumailleau *et al.*, 2009). In budding yeast, repolarization of the actin cytoskeleton during cytokinesis changes the itinerary of cargos from the growing bud to the bud neck and may play a role in the trafficking of daughter-specific cargos vs mother-specific cargos. We hypothesize differences in trafficking between mother and daughter may exist during cell separation, ensuring secretion of daughter-specific Ace2 targets are not inappropriately secreted from the mother.

Additionally, the evolutionarily conserved septins have been shown to control daughter-specific cell polarity in budding yeast (Barral *et al.*, 2000), and cells lacking septins fail to appropriately undergo cytokinesis as exocytic factors fail to localize properly (Caudron and Barral, 2009). Interestingly, several septin components have been shown to undergo asymmetric modification by SUMOylation (Johnson and Blobel, 1999; Takahashi *et al.*, 2001). An intriguing hypothesis is that this asymmetry informs which cargos and/or exocytic targeting factors become active during cell separation, leading to targeted exocytosis of specific cargos from the daughter.

Clearly *ACE2^3D^* mother cells produce Cts1, but it is unclear where Cts1 made in mother cells is trafficked. We favor a model in which diversion to the vacuole in mother cells could prevent secretion. Cts1 activity has been detected in isolated vacuoles in wild type cells (Elango *et al.*, 1982), and we find an increase in total Cts1 in wild type cells upon deletion of the major vacuolar protease *PEP4* (data not shown). This suggests that at least some of the intracellular Cts1 is shuttled to the vacuole, even in wild type cells. This model predicts that mother cells have a mechanism to target particular cargos to the vacuole. Alternatively, cargos may be trafficked to the vacuole by default until activation of an Ace2-independent daughter-specific factors re-direct vacuole-bound vesicles to the plasma membrane for secretion. Examination of these possibilities is an ongoing line of investigation.

Our study has begun to uncover the complexities of Cts1 regulation and suggests an asymmetry in how Cts1 is trafficked: secretion in daughters and degradation in mothers. This study suggest that a combination of factors control specification of the mother and daughter cell states, however the identity of these factors remain unknown. We recently demonstrated that the daughter-specific NDR/LATS kinase Cbk1 plays an Ace2-independent role in promoting Cts1 secretion (Brace *et al.*, 2018). However, the targets of Cbk1 that might control Cts1 secretion remain elusive. Taken together, while Ace2 helps to define the daughter cell state, our study demonstrates that Ace2-target asymmetry is not sufficient to induce the daughter cell state and additional, yet unidentified, intrinsic factors are required to reinforce the daughter cell identity.

## Materials and Methods

### Strains, plasmids, and growth conditions

All strains are derived from the W303 genetic background (*leu2-3,112 trp1-1 can1-100 ura3-1 ade2-1 his3-11,15 ssd1-d*). We used standard lithium acetate transformation and genetic crosses to isolate strains of the indicated genotypes in Table 1. The following methods were used to generate individual genotypes. We integrated the *ACE2*^3D^ (S122D, S137D and T436D) allele using a two-fragment PCR method to replace *ace2*D::*HIS3* with the mutant or wild type sequence, a GFP or HA tag and the KanMX marker. The *CTS1* and *VPS10* alleles were generated using homologous recombination replacement of a CORE cassette integrated into the genome using the Delitto Perfetto method (Storici and Resnick, 2006). For internal msGFP Cts1 tagging (imsNGFP-Cts1), three PCR reactions corresponding to Cts1 (1- 315), msGFP and Cts1 (316-562) were transformed into the *CTS1* CORE strain (ELY). Each PCR containing 20-30 base pair overlap between each other and with the genome for recombination. The monomeric-superfolder GFP (msGFP) sequence was amplified from a construct provided by Dr. Benjamin Glick (University of Chicago). Upon sequencing, we identified a single isolate (referred to as Cts1^trunc^) containing a serendipitous nucleotide deletion generating a truncation after the msGFP adding the following non-native sequence:LLQLPPQKPQQPQLHLLQLHLLQLLRKRPHNLRHLHKVKAKLLYLQLQAALSKHQLLKLQKH*. The Cts1 ΔCBD (chitin binding domain) was generated by CORE replacement with two PCR products corresponding to Cts1 (1-480) and msGFP. In all constructs, we sequenced to confirm proper integration. In all strains the endogenous 5’ and 3’ sequences remain intact and PCR was used for genotyping purposes in genetic crosses. To generate the *vps10-104* allele (Jørgensen *et al.*, 1999), we replaced a CORE integrated at *VPS10* to delete domain 1. The resulting unmarked locus was sequenced and PCR was used for subsequent genotyping in genetic crosses. We tagged Dse1 C-terminally with GFP using the Longtine method (Longtine *et al.*, 1998). The PLC-delta PH domain-mCherry (pYL95, provided by Scott Emr) was transformed into the indicated strains, and the MEP strain (DLY174, provided by Dr. Daniel Gottschling) was transformed with plasmids expressing *ACE2*-GFP (pELW755), *ACE2^3D^*-GFP (pELW2029) containing its endogenous 5’ and 3’ sequences or an empty vector (pELW69).

**Table 1:** Strains used in this study.

| Strain | Genotype | Source |
| --- | --- | --- |
| ELY83 | *leu2-3,112 trp1-1 can1-100 ura3-1 ade2-1 his3-11,15* (W303) | D. Drubin |
| ELY3670 | W303 NatNT2 :: GalL-*CDC20 ACE2*-GFP :: KanMX6 iHA-*CTS1* |  |
| ELY3595 | W303 NatNT2 :: GalL-*CDC20 ACE2<sup>3D</sup>* (122D, 137D, 436D) - GFP :: KanMX6 iHA-*CTS1* |  |
| ELY871 | *ACE2* - GFP :: KanMX6 |  |
| ELY3674 | *ACE2<sup>3D</sup>* (122D, 137D, 436D) - GFP :: KanMX6 |  |
| ELY3588 | *his3Δ1 leu2Δ0 ura3Δ0 lys2Δ0 HOΔ::SCW11Pr-Cre-EBD78-NATMX loxP-CDC20-intron-loxP::HPHMX loxP-UBC9-loxP::LEU2* + [pELW69] | D. Gottschling |
| ELY3589 | *his3Δ1 leu2Δ0 ura3Δ0 lys2Δ0 HOΔ::SCW11Pr-Cre-EBD78-NATMX loxP-CDC20-intron-loxP::HPHMX loxP-UBC9-loxP::LEU2* + [pELW755 *ACE2*] |  |
| ELY3681 | *his3Δ1 leu2Δ0 ura3Δ0 lys2Δ0 HOΔ::SCW11Pr-Cre-EBD78-NATMX loxP-CDC20-intron-loxP::HPHMX loxP-UBC9-loxP::LEU2* + [pELW2029 *ACE2<sup>3D</sup>*] |  |
| ELY3864 | *ACE2* - 3HA :: KanMX6 *imsNGFP* – *Cts1 vps10-104* |  |
| ELY3865 | *ACE2<sup>3D</sup>* (122D, 137D, 436D) - 3HA :: KanMX6 *imsNGFP* – *Cts1 vps10-104* |  |
| ELY3845 | *ACE2* - 3HA :: KanMX6 *Dse1* – GFP :: His3MX6 |  |
| ELY3846 | *ACE2<sup>3D</sup>* (122D, 137D, 436D) - 3HA :: KanMX6 *Dse1* – GFP :: His3MX6 |  |
| ELY3908 | *ACE2* - 3HA :: KanMX6 *Cts1 Δ-CBD msGFP vps10-104* + [pYL95: PLC-delta PH domain – mCherry :: Leu] |  |
| ELY3909 | *ACE2<sup>3D</sup>* (122D, 137D, 436D) - 3HA :: KanMX6 *Cts1 Δ-CBD msGFP vps10-104* + [pYL95: PLC-delta PH domain – mCherry :: Leu] |  |

We cultured cells in either YPD medium (1% yeast extract (BD), 2% peptone (BD), and 2% glucose (EMD)) or for microscopy we used synthetic minimal medium (SD) (0.67% yeast nitrogen base without amino acids (US Biological), 0.2% amino acid drop-in (US Biological), adenine (Ameresco), and 2% glucose (EMD)). In figure 1 and 2A, to examine asynchronous cells, ELY3670 and ELY3595 were grown in media containing 2% galactose (SC + Gal) to induce the *CDC20* gene and maintain cell cycle progression. To visualize external Cts1, synthetic minimal media was buffered to ~pH 7.3 with 10 mM Tris pH 8.4. Otherwise, unbuffered acidic media (~pH 4.0) was used. We cultured cells at 25° C unless otherwise indicated in the figure legend.

### Mother staining

An overnight culture grown to 1-2 ODs in SD or SC + Gal media was incubated for 10 minutes at room temperature with rhodamine-concanavalin A (Vector Laboratories) at a final concentration of 40ug/ml. Cells were washed twice in fresh media, diluted to 0.1-0.2 ODs into fresh media and grown at 25° C for 3-4 hours before imaging with a Texas Red compatible filter set.

### Cell separation quantification

Cells were grown to mid-log phase, sonicated for 30 seconds in a water bath sonicator, and imaged on an Axiovert 200 M microscope (Carl Zeiss MicroImaging, Inc.) fit with a 100×/1.45- numerical aperture oil immersion objective and Cascade II-512B camera (PhotoMetrics, Inc.). The number of connected cells in a clump was counted for three independent trials with >120 clumps per trial and genotype. The pooled data (*ace2Δ* n=382 groups, *ACE2* n=814 groups, *ACE2*^3D^ n=788 groups) was plotted as a box and whisker (5 to 95 percentile) plot in GraphPad Prism version 5.03. The “+” indicates the population mean.

### Bud scar stain

Mother cells were stained as above and grown to mid-log phase in SD media and then stained with a final concentration of 100μg/ml calcofluor (Fluorescent brightener 28; Sigma) for 10 minutes in the dark at 24°C. Cells were then washed twice and imaged with a DAPI compatible filter set. For bud scar quantification, we analyzed maximum projections of 0.2 um z-stacks and counted the number of bud scars and the presence or absence of rhodamine-ConA signal. Cells were binned into 1, 2-3 or 4+ bud scars. The number of bud scars per mother or daughter was counted in three independent experiments (*ACE2* n=46, 28, 45 and *ACE2*^3D^ n=57, 30, 39). A two-tailed, equal variance student’s t-test was determined using EXCEL to find there was no significant difference between *ACE2* and *ACE2*^3D^ (p > 0.05) for each bud scar category. *ACE2* mother vs *ACE2*^3D^ mother: one bud scar p=0.265, two to three bud scars p=0.307, and four or more bud scars p=0.326. *ACE2* daughter vs *ACE2*^3D^ daughter: one bud scar p=0.876 and two to three bud scars p=0.374.

### Microscopy

In figures 1D, 2A and S1 images were acquired with an Axiovert 200 M microscope (Carl Zeiss MicroImaging, Inc.) fit with a 100×/1.45- numerical aperture oil immersion objective and Cascade II-512B camera (PhotoMetrics, Inc.). A z-series of 0.2 μ m step size was taken using Openlab software (v5.5.0; Improvision), and FIJI and Photoshop (Adobe) were used to make linear adjustments to brightness and contrast. In figures 4B, 4D, 5A and S2B, images were acquired with a DeltaVision Core fit with a U PLAN S APO 100×/1.4 NA objective (Olympus) and a CoolSnapHQ2 Camera (Photometrics). A z-series of 0.2 μ m step size was taken. Images were deconvolved using softWoRx’s (Applied Precision Ltd.) iterative, constrained three-dimensional deconvolution method. FIJI was used to make linear adjustments to brightness and contrast. A single slice or the maximum projection, as indicated in the figure legend, is shown.

### Image Quantification

For Ace2 localization quantification, large-budded, mother-daughter pairs were identified and if mean fluorescent intensity of the nucleus was greater than the cytoplasmic mean, the pair was scored as ‘daughter nucleus’, ‘mother nucleus’ or ‘mother-daughter nucleus’. For intensity measurements, we drew a circular region of interest (ROI) around each nucleus. In cells without nuclear Ace2, a similar sized circle was drawn in the cytoplasm of the corresponding cell. Signal intensity was measured in each slice of the z-stack and the slice with maximum signal was determined. We then calculated the corrected intensity by subtraction of the mean background signal from that slice times the area of the ROI. We divided the intensity by 10,000 before graphing (arbitrary units). For Cts1 localization quantification, the analysis was limited to large-budded, mother-daughter pairs. Using FIJI threshold and particle analysis functions, ROI around cells expressing GFP signal over background were determined. The location of the ROI was then determined to be ‘daughter’, ‘mother’ or ‘mother-daughter’. For Dse1 localization quantification, the analysis was limited to large-budded, mother-daughter pairs. Cells were scored as ‘daughter’, ‘mother’ or ‘mother-daughter’ localized. All graphs were generated using GraphPad Prism version 5.03. To determine significance between the *ACE2* and *ACE2^3D^* mother and daughter localization of Cts1 and Dse1 a two-tailed, equal variance student’s t-test was calculated in EXCEL.

For Cts1 secretion analysis, in FIJI a 2-micron line was drawn across the bud neck at the secretion site from mother to daughter cell in a single slice exhibiting clear separation of the mother-daughter plasma membrane (PM) signal. The line’s center was aligned between the mother and daughter PM and marked zero. Signal intensity across the line was measured and transferred to Excel. The green and red channel intensities were normalized by scaling data from 0 to 1 using the equation x’ = (x-min(x)) / (max(x) – min (x)). The average signal intensity with standard error of the mean was calculated and graphed in GraphPad Prism version 5.03. The distance between the Cts1 and plasma membrane peaks was determined by subtracting the distance at which the peak Cts1 occurred from each PM. In one case, the Cts1 signal was not between the PM peaks giving a negative value. Mother cells were identified by either size difference (mothers being larger) or the presence of one or more bud scars upon calcofluor staining. To determine the significance of signal distance between each genotype and cell type a two-tailed, unpaired student’s t-test was calculated in GraphPad Prism version 5.03. *ACE2* daughter vs *ACE2*^3D^ daughter p=0.914, *ACE2* mother vs *ACE2*^3D^ mother p=0.497, *ACE2* daughter vs *ACE2* mother p =0.016, *ACE2*^3D^ daughter vs *ACE2*^3D^ mother p=0.040, *ACE2* daughter vs *ACE2*^3D^ mother p=0.057.

### Mother Enrichment Program spot assays

Cells were grown to mid-log and then plated in five-fold serial dilutions on plates containing 1μM estradiol (MEP active) or the equivalent volume of ethanol (MEP inactive). Cells were incubated at 24° C for 3 days, then replica plated to plates containing ethanol. Cells were permitted to grow for an additional 2 days at 24°C.

### RNA preparation and qPCR

We prepared RNA from *ACE2* or *ACE2^3D^* asynchronous cells by hot acid phenol extraction as previously described (Collart and Oliviero, 1993). We treated 2 mg of RNA with 10 units of RNase-free DNase I (Roche) and converted it to cDNA with Moloney murine leukemia virus reverse transcriptase (MMLV-RT; Promega). We performed quantitative RT-PCRs (qPCR) with the iCycler Thermal Cycler with iQ5 Multicolor Real-Time PCR Detection System (Bio-Rad) using primers specific to *CTS1, SCW11, DSE1, DSE2, DSE3, DSE4*, and *ACT1*. We generated standard curves from serial dilutions of intact yeast genomic DNA using linear regression analysis of cycle threshold (CT) values. Samples were internally normalized to the CT of *ACT1* and the fold change relative to the control sample (*ACE2*) is shown.

### Western blot

One O.D. of cells grown to mid-log phase were harvested and resuspended in 1mL 0.255 M NaOH + 0.1% BME and placed on ice for 10 minutes. 138μL of 50% TCA was added and cells were left on ice for another 10 minutes. Cells were pelleted at 4°C at 13,000rpm for 10 minutes in a microcentrifuge. Pellets were washed with 1mL cold acetone and the resulting pellet was dried and resuspended in 40μL of MURB (200mM MES Buffer pH 7, 2% SDS, 6 M Urea). 15μL of lysate was loaded on a 10% SDS-PAGE gel and transferred to Immobilon FL PVDF (polyvinylidene difluoride; Millipore) membrane. The membrane was blocked for 30 minutes in Odyssey blocking buffer PBS (Licor), then incubated with primary antibody in Tris-buffered saline (50 mM Tris pH 7.6, 150 mM NaCl) plus 0.1% Tween (TBS-T) for 1 h at room temperature. We then incubated blots with secondary antibody in TBS-T plus 0.1% SDS for 30 minutes. Membranes were washed three times for 3 minutes with TBS-T between antibody additions and prior to imaging. We used primary antibodies as follows: mouse monoclonal Pgk1 (22C5D8, ThermoFisher) at 1:1000, rabbit polyclonal GFP (A-11122, ThermoFisher) at 1:2000. We used secondary antibodies as follows: IRDye 680 LT goat anti-mouse (Licor) at 1:20,000 or IRDye 800 CW goat antirabbit (Licor) at 1:15,000. We imaged and processed blots with the Image Studio Lite Odyssey software (v4.0; Li-Cor).

## Acknowledgments

We thank members of the Weiss lab, Laura Lackner, and members of her laboratory at Northwestern University for critical review of the manuscript and invaluable discussions. We thank Dr. Benjamin Glick and Dr. Daniel Gottschling for strains and plasmids. We also thank staff and instrumentation support from the Biological Imaging Facility and High Throughput Analysis Laboratory at Northwestern University. Research reported in this publication was supported by a grant awarded to E.L.W. from the National Institutes of Health (NIH) under award number R01GM084223.

## Author Contributions

Conceptualization, V.N.T, J.L.B and E.L.W; Methodology, V.N.T, J.L.B and E.L.W; Investigation, V.N.T and J.L.B; Writing, V.N.T, J.L.B and E.L.W; Visualization, V.N.T, J.L.B, and E.L.W; Funding Acquisition, E.L.W.

## Supplemental Figures

**Figure S1.**
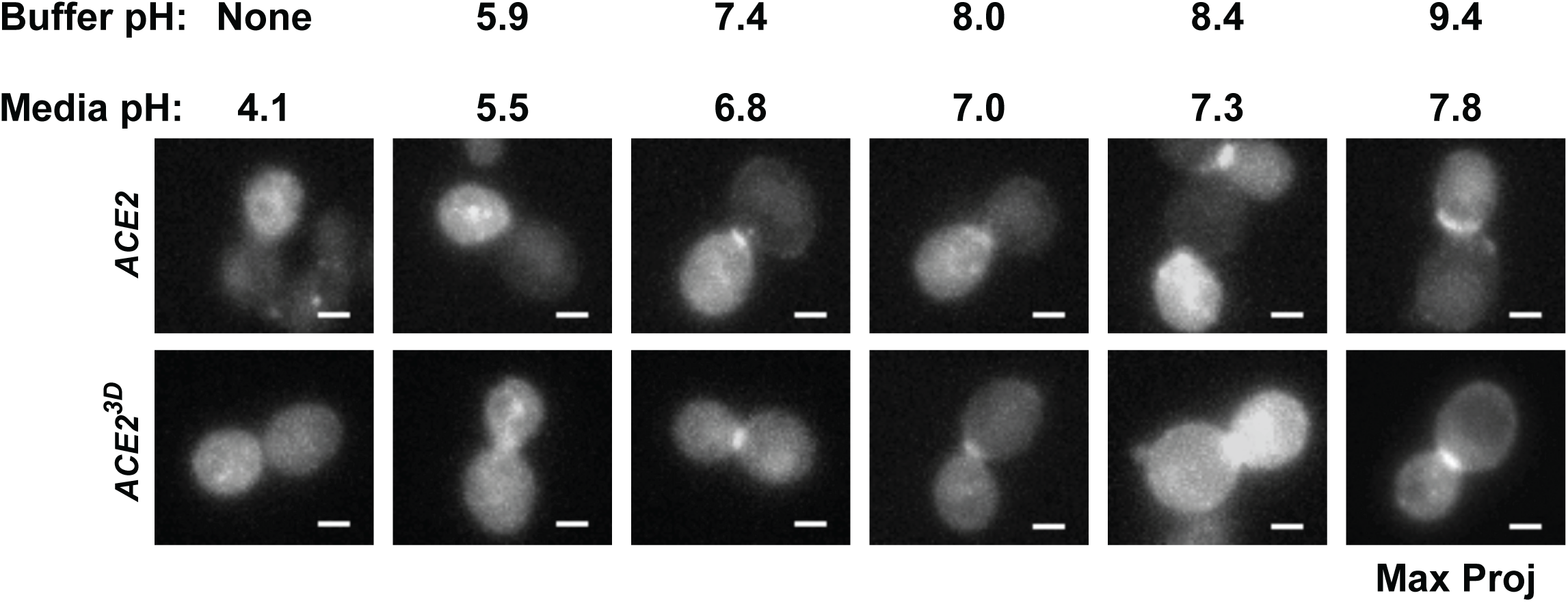
Related to Figure 4. Effect of pH on msGFP-Cts1 fluorescence signal. Representative maximum projection images of *ACE2 vps10-104* or *ACE2^3D^ vps10-104* cells expressing Cts1-msGFP grown to log phase. A single population of cells were washed and resuspended in minimal media buffered to the indicated pH before imaging. Scale bar is 2 microns.

**Figure S2.**
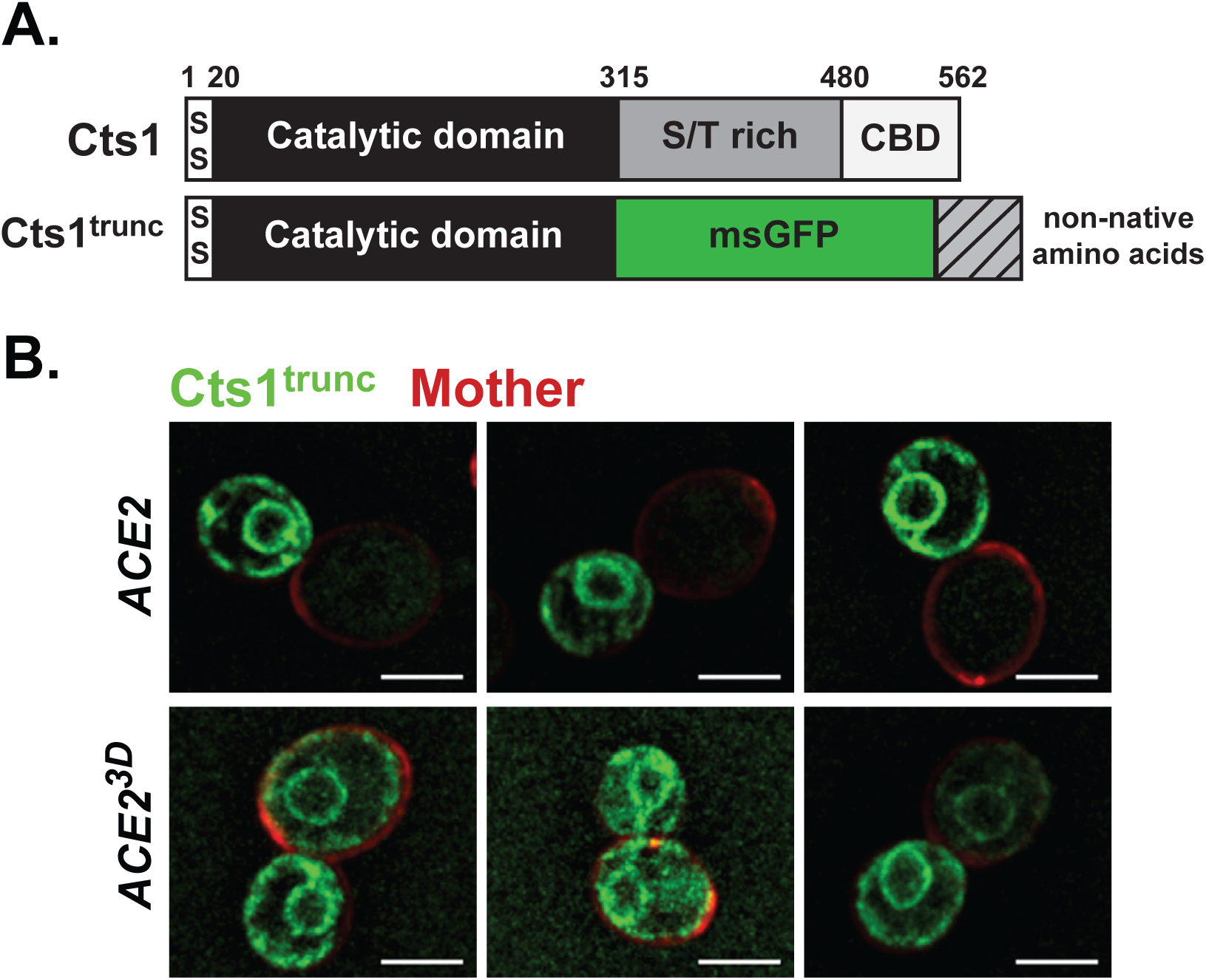
Related to Figure 4. Schematic of Cts1 and the truncation allele (Cts1trunc). A serendiptious deletion created a premature stop codon after the msGFP shifting the reading frame so that additional non-native amino acids are coded for at the C-terminus of Cts1 (see methods). Representative single slice images of *ACE2* and *ACE2^3D^* cells expressing Cts1trunc (green). Mother cells are stained with rhodamine-ConA (red). Scale bar is 5 microns.

## Abbreviations

ER: Endoplasmic reticulum
ConA: Concanavalin A
MEP: Mother Enrichment Program
msGFP: Monomeric superfolder GFP
CBD: Chitin Binding Domain
PM: Plasma membrane

